# The Regulator FleQ Post-Transcriptionally Regulates the Production of RTX Adhesins by *Pseudomonas fluorescens*

**DOI:** 10.1101/2023.05.09.540025

**Authors:** Alexander B. Pastora, George A. O’Toole

## Abstract

Biofilm formation by the Gram-negative gammaproteobacterium *Pseudomonas fluorescens* relies on the production of the repeat-in-toxin (RTX) adhesins LapA and MapA in the cytoplasm, secretion of these adhesins through their respective type 1 secretion systems, and retention at the cell surface. Published work has shown that retention of the adhesins occurs via a post-translational mechanism involving the cyclic-di-GMP receptor LapD and the protease LapG. However, little is known about the underlying mechanisms that regulate the production of these adhesins. Here, we demonstrate that the master regulator FleQ modulates biofilm formation by post-transcriptionally regulating the production of LapA and MapA. We find that a Δ*fleQ* mutant has a biofilm formation defect compared to the WT strain, which is attributed in part to a decrease in LapA and MapA production, despite the Δ*fleQ* mutant having increased levels of *lapA* and *mapA* transcripts compared to the WT strain. Through transposon mutagenesis and subsequent genetic analysis, we found that over-stimulation of the Gac/Rsm pathway partially rescues biofilm formation in the Δ*fleQ* mutant background. Collectively, these findings provide evidence that FleQ regulates biofilm formation by post-transcriptionally regulating the production of LapA and MapA, and that activation of the Gac/Rsm pathway can enhance biofilm formation by *P. fluorescens*.

**Importance:** Biofilm formation is a highly coordinated process that bacteria undergo to colonize a variety of surfaces. For *Pseudomonas fluorescens*, biofilm formation requires the production and localization of RTX adhesins to the cell surface. To date, little is known about the underlying mechanisms that regulate biofilm formation by *P. fluorescens*. Here, we identify FleQ as a key regulator of biofilm formation that modulates the production of LapA and MapA through a post-transcriptional mechanism. We provide further evidence implicating activation of the Gac/Rsm system in FleQ-dependent regulation of biofilm formation. Together, our findings uncover evidence for a mechanism of post-transcriptional regulation of the LapA/MapA adhesins.

## Introduction

*Pseudomonas fluorescens* is a Gram-negative gammaproteobacterium that is broadly distributed in the environment, and is commonly thought of as a plant-commensal bacterium (1). *P. fluorescens* colonizes plant roots and survives on nutrients secreted by these plants while producing antimicrobial and antifungal metabolites that are secreted into the surrounding rhizosphere (2–4). While commonly associated with the environment, *P. fluorescens* has become a clinically relevant organism in that it has been associated with cases of hospital acquired bacteremia via the contamination of hospital products (5–9) and has been associated with Crohn’s Disease whereby *P. fluorescens* colonizes the intestines and increases the permeability of intestinal epithelial cells (10–14). To persist in these diverse environments, *P. fluorescens* relies on the highly coordinated transition from a motile to biofilm lifestyle (15, 16)

For *P. fluorescens*, biofilm formation is predominantly dependent on the repeat-in-toxin (RTX) adhesins LapA and MapA, which mediate surface attachment and contribute to biofilm initiation and maturation when localized to the cell surface (17, 18). Localization of the adhesins to the cell surface is mediated by each adhesin’s respective Type 1 Secretion Systems (T1SS), although recent evidence suggests that the Lap system secretes MapA in the absence of LapA (18). Cell surface retention of the adhesins is mediated by the cyclic-di-GMP sensing protein LapD, which sequesters the periplasmic protease LapG when levels of cyclic-di-GMP are high. When cyclic-di-GMP levels are depleted, LapG is free to cleave the N-terminus of LapA and MapA at their characteristic TAAG motif, which releases these adhesins from the cell surface where they no longer contribute to biofilm formation (19). Additionally, previous work has shown that levels of cyclic-di-GMP can be altered in response to specific metabolites, such as inorganic phosphate and citrate, via stimulation of specific diguanylate cyclases and phosphodiesterases, which in turn affects LapD-dependent adhesin retention (20–23).

Apart from LapD-dependent regulation of adhesin retention, additional regulatory mechanisms affecting adhesin production have been poorly characterized for *P. fluorescens*. The conserved transcriptional regulator FleQ has been shown to regulate a variety of targets related to biofilm formation, such as motility and exopolysaccharide production, in multiple *Pseudomonas* spp. (24–29). FleQ has been associated with motility of *P. fluorescens* and there is some evidence that FleQ may regulate *lapA* gene expression (29, 30). Here, we demonstrate that FleQ post-transcriptionally regulates biofilm formation by modulating levels of LapA and MapA and provide evidence that this regulation occurs through the Gac/Rsm pathway.

## Results

### FleQ regulates biofilm formation through the post-transcriptional regulation of *lapA* and *mapA*

To assess the role of FleQ in biofilm formation, we made a chromosomal deletion of the *fleQ* gene in *P. fluorescens* Pf0-1 and probed the mutant for biofilm formation in a static biofilm assay using KA medium, a minimal medium supplemented with arginine, which has previously been shown to support the formation of robust, LapA- and MapA-dependent biofilms (18). After 16h of growth, the Δ*fleQ* mutant showed a significant decrease in biofilm formation compared to the wild-type (WT) strain. When the *fleQ* deletion mutation was complemented at the *att* site with the wild-type *fleQ* gene, biofilm formation was restored to levels similar to the wild-type (WT) strain (Fig. 1A). We additionally grew the WT and Δ*fleQ* mutant in a microfluidics device with continuous irrigation of KA medium and imaged the biofilm after five days of growth at room temperature. As with the static biofilm assay, the Δ*fleQ* mutant similarly has a biofilm formation defect under flow (Fig. S1).

**FIG 1.**
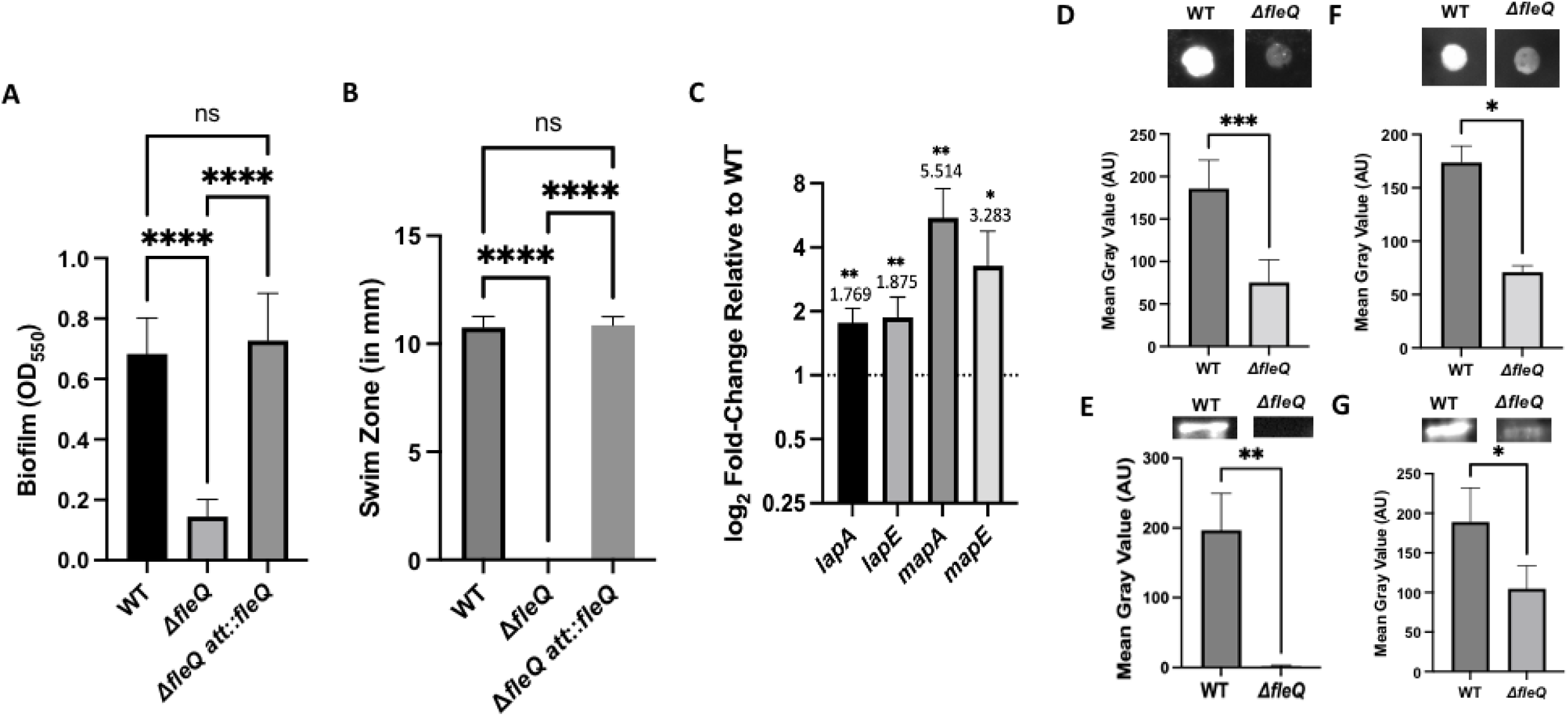
A FleQ-deficient strain has a biofilm formation defect due to a decrease in adhesin production. (A) Quantification of the biofilm formed by the WT strain, Δ*fleQ* mutant, and a Δ*fleQ* mutant complemented with the wild-type *fleQ* gene at the *att* site measured at OD_550_ after 16h of growth in KA minimal medium. Statistical significance was determined using an unpaired t-test. ****, P<0.0001. (B) Swim zone (in millimeters) of the WT strain, Δ*fleQ* mutant, and a Δ*fleQ* mutant complemented with the wild-type *fleQ* gene at the *att* site after toothpick inoculation on KA medium supplemented with 0.3% agar after 24h of growth at 30°C. Statistical significance was determined using an unpaired t-test. ****, P<0.0001. (C) Expression of *lapA*, *lapE*, *mapA*, and *mapE* genes in the Δ*fleQ* mutant relative to the WT strain, after 16h of growth on KA medium supplemented with 1.5% agar (KA agar), using the 2^-ΔΔCt^ (Livak) method. For statistical significance, paired t-tests were conducted for each gene between the WT strain and Δ*fleQ* mutant. *, P<0.05; **, P<0.01. (D,F) Quantification of cell surface-associated LapA after 16h of growth on KA agar (D) or MapA after 24h of growth on KA agar (F) for the WT strain and Δ*fleQ* mutant using ImageJ by measuring the mean gray value of each spot using a pre-defined region of interest (ROI) and subtracting the background, which was determined by measuring the mean gray value of a section of the blot without any sample. Representative images are included above each graph. Statistical significance was determined using unpaired t-tests. *, P<0.05; ***, P< 0.001. (E,G) Quantification of LapA after 16h of growth on KA agar (E) or MapA after 24h of growth on KA medium (G) from whole cell lysates that were prepared from 25 ml of cultures, concentrated to 100 µl in 3mg/mL lysozyme with sonication and quantified for total protein using the BCA assay. 25µg (E) or 50µg (G) of total protein was resolved on a 7.5% TGX Gel and then blotted for LapA or MapA, respectively. Representative images are included above each graph. Statistical significance was determined using unpaired t-tests. *, P<0.05; **, P<0.01. All error bars represent standard deviation.

Previous work demonstrated that a *P. fluorescens* Pf0-1 transposon insertion mutant in FleQ is non-motile and that FleQ regulates motility through the transcriptional regulation of a subset of the flagellar assembly genes in *Pseudomonas ogarae* F113, a recently reclassified strain of *P. fluorescens* (30–33). We thus assessed motility of the Δ*fleQ* mutant via swim assay using KA medium supplemented with 0.3% agar, which revealed that this mutant is unable to swim. When the *fleQ* detetion mutant was complemented at the *att* site with the wild-type *fleQ* gene, swimming motility was restored to levels similar to the wild-type (WT) strain (Fig. 1B).

Previous work showed that biofilm formation by *P. fluorescens* Pf0-1 is dependent on LapA and MapA production and subsequent localization to the cell surface (17, 18). To determine the effect(s) of FleQ on production of these adhesins, we first assessed the expression levels of the genes encoding LapA and MapA, as well as the genes encoding their respective outer membrane porins LapE and MapE, in the WT and Δ*fleQ* mutant grown on KA agar. Interestingly, the Δ*fleQ* mutant shows modest (∼1.8-5-fold), but significant increase in expression of all four genes compared to the WT strain (Fig. 1C).

We next assessed whether LapA or MapA protein production were negatively impacted by the absence of FleQ. To assess the impact on the LapA and MapA proteins, we probed for these adhesins on the cell surface using dot blot analysis and at the whole cell level using whole cell lysate Western blot, as described in the Materials and Methods. After 16h of growth on KA agar, LapA is present at the cell surface in the WT and readily detected in whole cell lysates, while the Δ*fleQ* mutant has no detectable LapA at the cell surface or in whole cell lysates (Fig. 1D-E).

Interestingly, MapA was not readily detected at the cell surface of the WT strain or the Δ*fleQ* mutant after 16h of growth (Fig. S2). However, when growth on KA agar was increased to 24h, MapA is detected at the cell surface in the WT strain, while MapA is decreased in the Δ*fleQ* mutant at the cell surface (Fig. 1F). Interestingly, despite scaling up the amount of cells harvested and increasing the protein concentration, levels of MapA in the WT strain were below the limit of detection in whole cell lysates when grown on KA agar at 24h. However, when cultures were grown in KA liquid medium for 24h in a volume of 25 ml and then concentrated to 100 µl as described in Materials and Methods, MapA is detected in whole cell lysate in the WT while this protein showed reduced levels in the Δ*fleQ* mutant (Fig. 1G).

### Genetic studies reveal that LapA and MapA are produced at reduced levels in a FleQ- deficient strain

While we were unable to readily detect the LapA or MapA proteins in the Δ*fleQ* mutant, we sought to determine if a basal amount of these adhesins is still produced in this mutant, and whether forcing these adhesins to the cell surface can restore biofilm formation in the Δ*fleQ* mutant. Since previous work has shown that loss of LapG in the WT strain leads to the retention of LapA and MapA on the cell surface (34–36), we first made a chromosomal deletion of the *lapG* gene in the Δ*fleQ* mutant and assessed the Δ*lapG*Δ*fleQ* double mutant for its ability to form a biofilm. After 24h of growth in KA medium, the Δ*fleQ* Δ*lapG* mutant shows a significant increase in biofilm formation compared to the Δ*fleQ* mutant (Fig. 2A), but the biofilm is still reduced compared to the WT.

**FIG 2.**
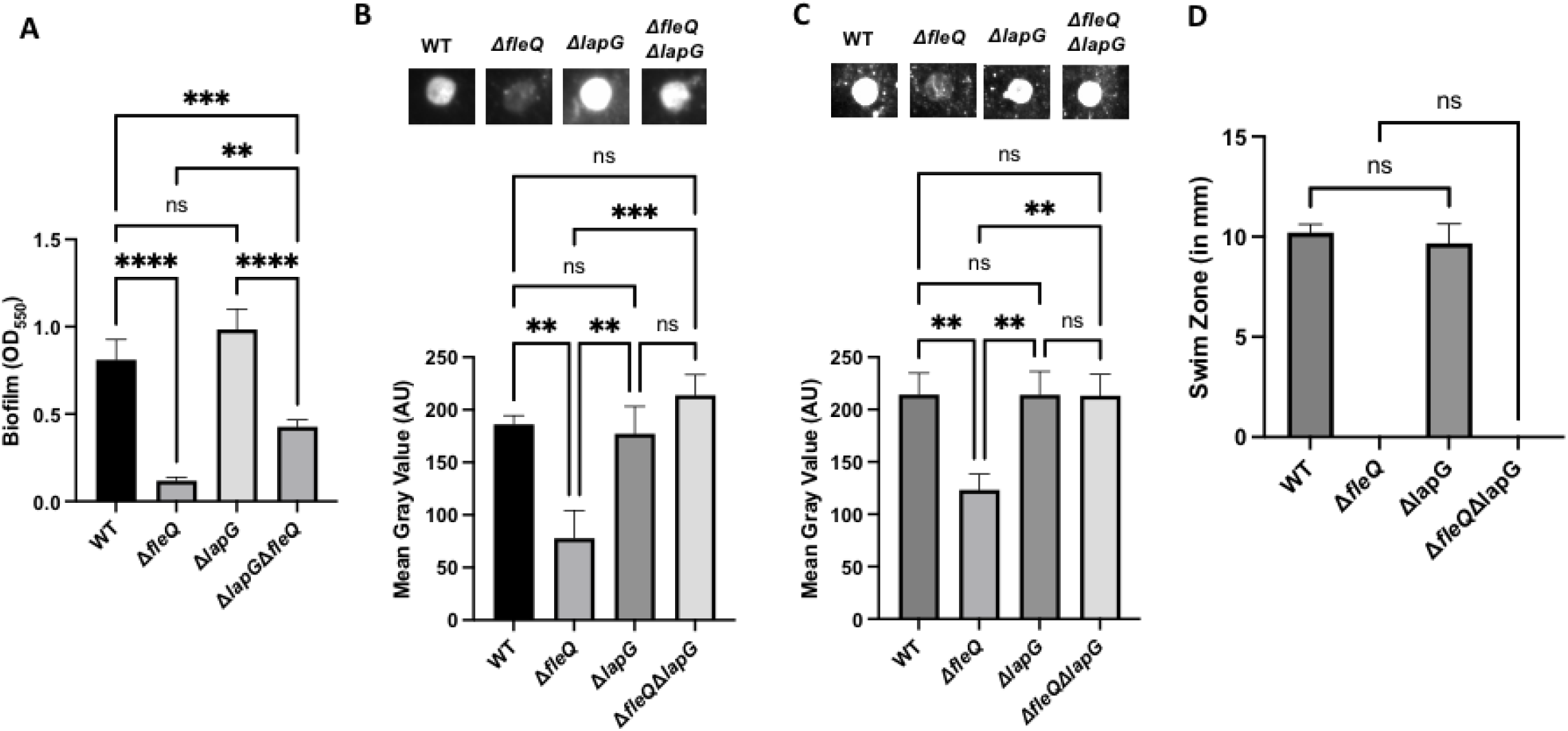
A FleQ-deficient strain produces adhesin sufficient to restore biofilm formation in a *lapG* mutant. (A) Quantification of the biofilm formed for the WT and the Δ*fleQ,* Δ*lapG*, and Δ*fleQ*Δ*lapG* mutant strains measured at OD_550_ after 24h of growth in KA minimal medium. This timepoint was chosen because both LapA and MapA are detected at the cell surface at this time point. (B-C) Quantification of cell surface-associated LapA after 16h of growth on KA agar (B) or MapA after 24h of growth on KA agar (C) in the WT, Δ*fleQ,* Δ*lapG*, and Δ*fleQ*Δ*lapG* strains as described in Figure 1. Representative images are included above each graph. (D) Swim Zone (in millimeters) for the WT, Δ*fleQ,* Δ*lapG*, and Δ*fleQ*Δ*lapG* strains after toothpick inoculation on KA supplemented with 0.3% agar and 24h of growth at 30°C. Statistical significance for this figure was determined using one-way ANOVAs with Tukey’s multiple comparisons tests. **, P<0.01; ***, P<0.001. All error bars represent standard deviation.

Given these findings, we then assessed the Δ*fleQ* Δ*lapG* mutant for surface associated LapA and MapA using dot blot analysis at 16h or 24h, respectively, as described in Figure 1. Here, we see that LapA and MapA are both detected at the cell surface in the Δ*fleQ* Δ*lapG* mutant but not in the Δ*fleQ* mutant (Fig. 2B-C). This result suggests that LapA and MapA are being produced, but below the limit-of-detection of the whole cell lysate Western blot in a FleQ-deficient strain. This finding indicates that these proteins can be detected if locked on the cell surface in the *lapG* mutant background (Fig, 2B-C). Consistent with this result, we find that over-expressing the LapBCE ABC transporter, required for LapA surface localization (17) and that we showed previously increases LapA surface localization (37), also partially rescues the biofilm defect of the *fleQ* mutant (Fig. S3).

While LapG-dependent regulation has not previously been associated with motility, we also assessed the Δ*fleQ* Δ*lapG* mutant for any impacts on motility using a swim assay. After 24h of growth, the Δ*fleQ* Δ*lapG* mutant, like the Δ*fleQ* mutant was unable to swim, while the WT strain and the Δ*lapG* mutant had similar swim zones (Fig. 2D).

### Mutational analysis of the *lapA* promoter reveals that FleQ transcriptionally regulates *lapA*

Previous work investigating FleQ-dependent gene regulation of *Pseudomonas aeruginosa* demonstrated that FleQ transcription occurs through the differential binding of the target promoter at two binding boxes. This work identified a binding box consensus sequence that was used to predict putative FleQ-regulated genes in other *Pseudomonas* spp. (28, 29). Interestingly, two FleQ binding boxes were identified within the predicted promoter region of the *lapA* gene of *P. fluorescens* Pf0-1 (29). To assess the function of these putative binding boxes, we mutagenized either Box 1 or Box 2 of the WT strain or the Δ*fleQ* mutant by altering the guanine in position 1 and cytosine in position 14 to adenine (Fig. 3A). We selected these nucleotides as they are the most conserved within the binding box consensus sequence (29).

**FIG 3.**
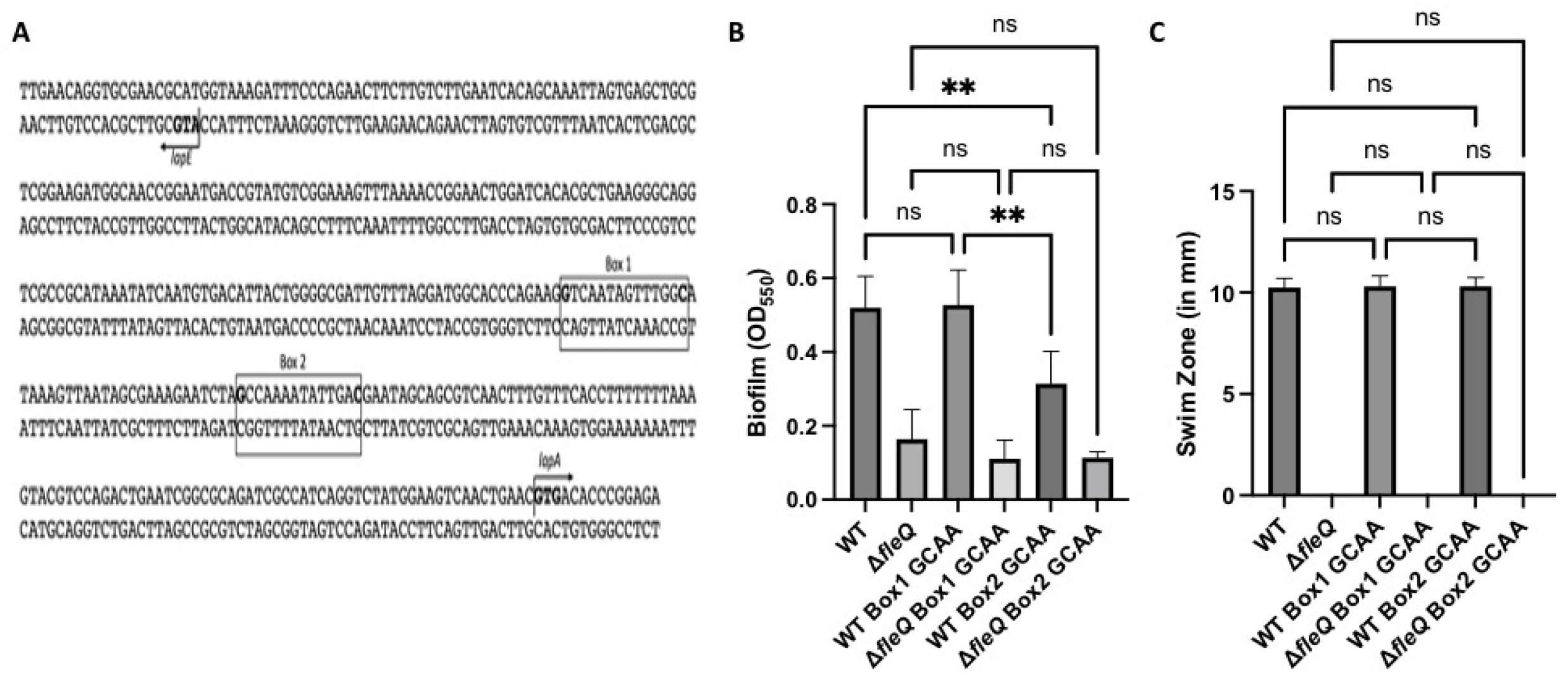
Mutagenizing the FleQ consensus sequences in the *lapA* promoter has a transcriptional effect that is masked in the Δ*fleQ* mutant. (A) Schematic showing the divergent *lapA*/*lapE* promoter. FleQ binding Box 1 and Box 2 are illustrated, and the respective sites of the point mutations are highlighted in bold. The *lapA* and *lapE* start codons are highlighted in bold and direction of translation is indicated by an arrow. (B) Quantification of the biofilm formed for the WT and the Δ*fleQ* mutant with a mutated FleQ binding Box 1 or Box 2, or the WT sequence at OD_550_ after 24h of growth in KA minimal medium. (C) Swim Zone (in millimeters) of WT and Δ*fleQ* strains with a mutated FleQ binding Box 1 or Box 2, or the WT sequence after toothpick inoculation on KA medium supplemented with 0.3% agar and 24h of growth at 30°C. Statistical significance for this figure was determined using one-way ANOVAs with Tukey’s multiple comparisons tests. **, P<0.01. All error bars represent standard deviation.

We then assessed these mutants for their ability to form a biofilm with the static biofilm assay. After 24h of growth in KA medium, the Box 1 mutant in the WT and Δ*fleQ* mutant backgrounds showed no significant change in biofilm formation compared to their parent strains. However, the Box 2 mutant in the WT background showed a significant decrease in biofilm formation compared to the WT strain. The Box 2 mutant in the Δ*fleQ* mutant background showed no significant change in biofilm formation as compared to the Δ*fleQ* parent strain (Fig. 3B), but it is important to note that the Δ*fleQ* mutant already makes a much-reduced biofilm compared to the WT. These data, along with the previously shown expression and protein data (Fig. 1D-E), suggest that FleQ may regulate *lapA* transcription via the Box 2 binding site.

We used the swim assay to verify that motility in these strains were unaffected. As expected, the *lapA* Box 1 and Box 2 mutants had similar swim zones to their parent strains (Fig. 3C), demonstrating that the changes in biofilm formation were not a consequence of reduced motility. These data also support the conclusion that loss of FleQ impacts biofilm formation by via reduced motility and reduced production of LapA.

### Transposon mutagenesis reveals additional factors that contribute to FleQ-dependent regulation of biofilm formation

The observations above that mutating the *fleQ* gene results in a modest (but significant) increase in *lapA* and *mapA* gene expression, but a large reduction in whole cell (and thus surface) levels of the LapA and MapA proteins, suggests that FleQ plays a positive, post-transcriptional role in the regulation of LapA and MapA proteins. Given the hypothesized post-transcriptional effect observed for the Δ*fleQ* mutant, we sought to identify additional factors that contribute to FleQ-dependent biofilm formation by performing a genetic screen that restores biofilm formation to the *fleQ* mutant. Briefly, the Δ*fleQ* mutant was conjugated as recipient with an *Escherichia coli* strain donor containing a plasmid harboring the Tn*M* mariner transposon, and Δ*fleQ* mutants with transposon insertion were selected by growth on LB agar supplemented with gentamycin (selection) and chloramphenicol (counter-selection). Individual insertion mutants were screened for their ability to form a biofilm and candidates with a significant increase in biofilm formation compared to the parent strain were chosen for further analysis.

The location of transposon insertion in these biofilm-forming candidate strains was determined via arbitrary primed PCR and sequencing of the flanking regions, as described in the Materials and Methods. From this screen, we identified four mutants that mapped to: the *lapG* gene coding sequence, the *flhA* gene coding sequence, intergenic region between the *flhA* and *flhF* genes, and the promoter region of the *gacS* gene (Fig. 4A). All four mutants showed a significant increase in biofilm formation compared to the Δ*fleQ* parental strain after 24h of growth on KA medium (Fig. 4B). We also assessed the motility of all four mutants; none of the four mutants were able to swim on KA supplemented with 0.3% agar (Fig. 4C), suggesting that the increase in biofilm formation was not related to changes in motility.

**FIG 4.**
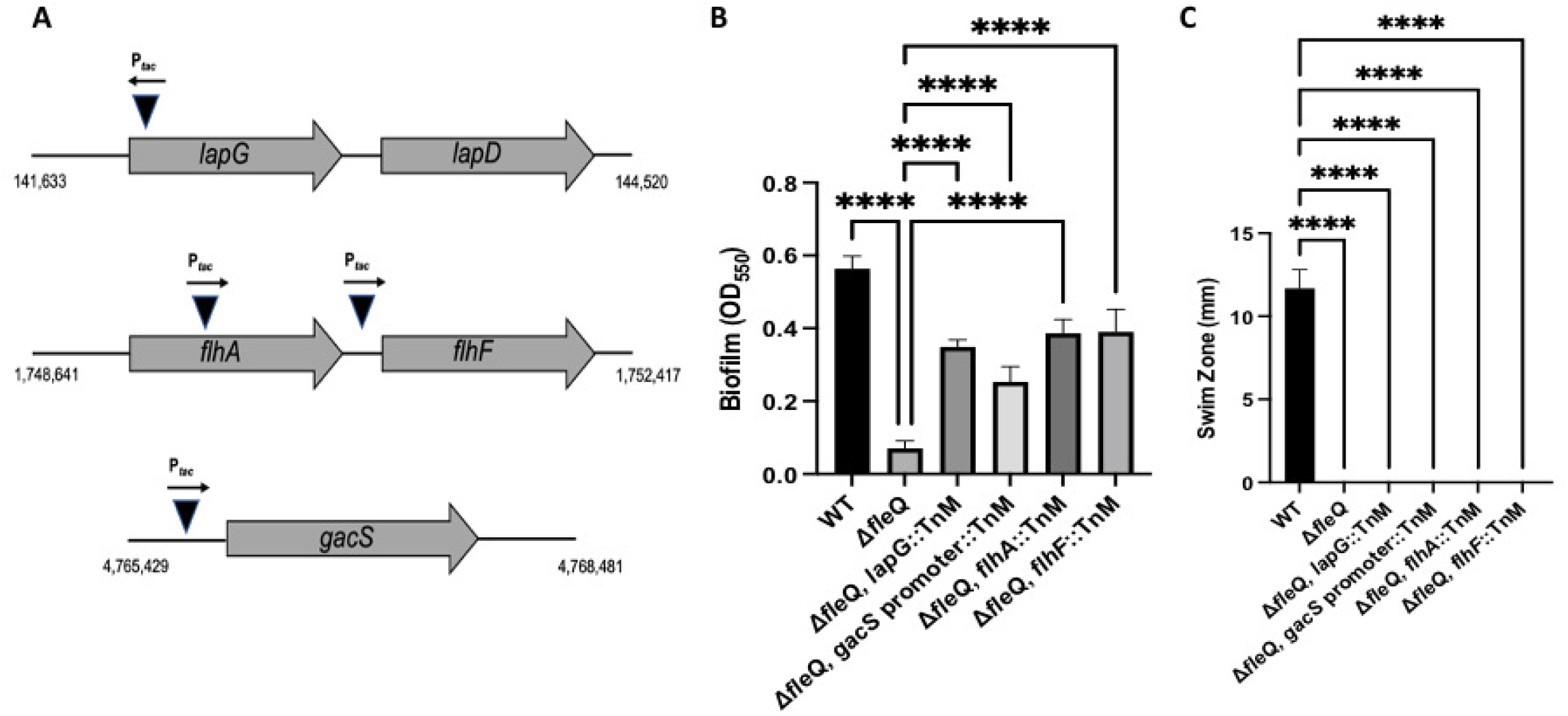
Identifying other factors that contribute to FleQ-dependent regulation of adhesins. (A) Schematic showing the location of FleQ-deficient transposon mutants that restore biofilm formation. Triangles indicate insertion into the genome and the arrow above each triangle indicates directionality of the P*tac* promoter. Genomic coordinates are included for reference. (B) Quantification of the biofilm formed by the WT strain, the Δ*fleQ* mutant and the indicated Δ*fleQ* derivatives at OD_550_ after 24h of growth in KA minimal medium. (C) Swim Zone (in millimeters) measurements of the WT strain, the Δ*fleQ* mutant, and Δ*fleQ* derivatives after toothpick inoculation on KA medium supplemented with 0.3% agar and 24h of growth at 30°C. Statistical significance for this figure was determined using one-way ANOVAs with Tukey’s multiple comparisons tests. ****, P<0.0001. All error bars represent standard deviation.

### Stimulation of the Gac/Rsm pathway rescues biofilm formation in a FleQ-deficient strain

Given our findings from the genetic screen, we hypothesized that the changes in biofilm formation for the four mutants identified were likely due to changes in adhesin production or their localization to the cell surface. For the *lapG* insertion mutant, we have already demonstrated that the loss of LapG in a Δ*fleQ* mutant leads to an increase in biofilm formation, which is due to increased localization of LapA and MapA to the cell surface (Fig. 2A-C). Thus, identifying a mutation in the *lapG* gene served to help validate the screening approach.

Our preliminary genetic analysis suggests that the *flhAF* operon may be involved in biofilm formation in a FleQ-independent manner (data not shown). Analysis of the *flhAF* operon and its role in biofilm formation will be addressed in a separate study.

We further analyzed the impact of the insertion mutation in the promoter of the *gacS* gene. First, given that the transposon carries a constitutive P*_tac_* promoter oriented in a manner that could drive expression of the *gacS* gene (Fig. 4A), we assessed the impact of the insertion on *gacS* gene expression through quantitative reverse transcription PCR. The insertion mutation in the promoter of the *gacS* gene results in expression of this gene to a level that is ∼20 times higher than the *gacS* gene is expressed in the Δ*fleQ* mutant or the WT strain (Fig. S4).

Previous work in *P. aeruginosa* and *Pseudomonas protogens* demonstrated that the Gac/Rsm pathway is regulated by the GacS-GacA two component system. The sensor kinase GacS phosphorylates its response regulator GacA, which then binds to the upstream activating sequence of its targets and upregulates transcription of small regulatory RNAs (38–43). These small regulatory RNAs are known to bind the RsmA, RsmE, and RsmI proteins (collectively, the “Rsm proteins”) and thereby prevent the Rsm proteins from binding to the mRNA of their target genes (41, 44–46).

Given the increased biofilm formation of the Δ*fleQ* mutant carrying an insertion mutation in the promoter of the *gacS* gene, and the increased level of *gacS* expression in this genetic background, we hypothesized that increased activation of the Gac/Rsm pathway might result in increased biofilm formation. To simulate constitutive overexpression of the Gac/Rsm pathway, we made chromosomal deletions of the *rsmA*, *rsmE*, and *rsmI* genes, which code for the three Rsm proteins, in both the WT strain and Δ*fleQ* mutant. We assessed the Δ*rsmA* Δ*rsmE* Δ*rsmI* triple mutant as well as individual deletion mutants for biofilm formation. After 24h of growth in KA medium, the Δ*rsmA* Δ*rsmE* Δ*rsmI* triple mutant shows a significant increase in biofilm formation in the Δ*fleQ* background (Fig. 5A). The Δ*rsmE* single mutant also shows a significant increase in the Δ*fleQ* background, whereas the Δ*rsmA* and Δ*rsmI* single mutants show no significant difference in biofilm formation in either strain background (Fig. S5A), suggesting that this phenotype may be largely RsmE-dependent.

**FIG 5.**
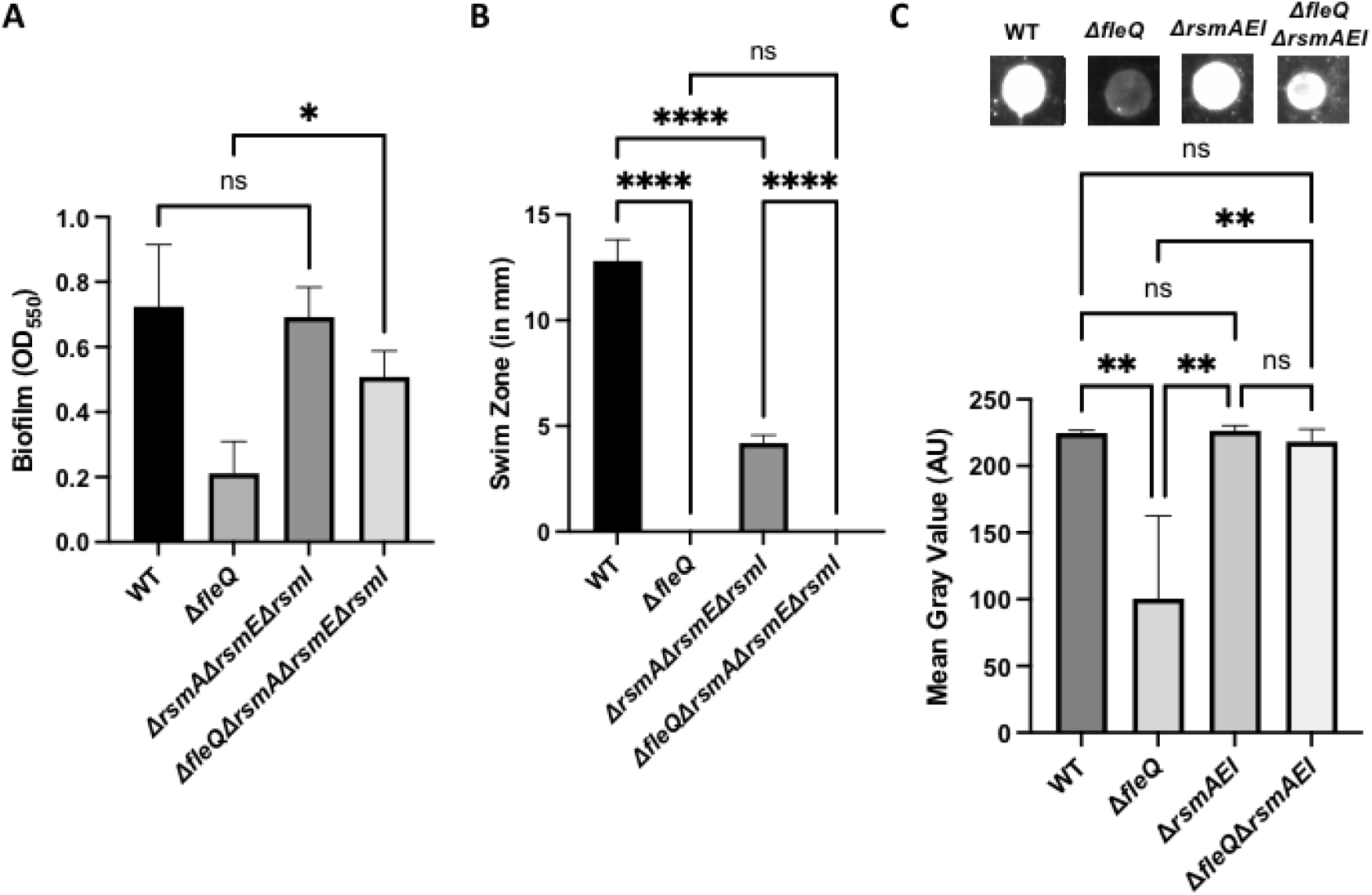
Modulation of the Gac/Rsm system increases biofilm formation in a WT and FleQ-deficient strain. (A) Quantification of the biofilm formed by the WT, Δ*fleQ,* Δ*rsmA*Δ*rsmE*Δ*rsmI*, and Δ*fleQ* Δ*rsmA*Δ*rsmE*Δ*rsmI* strains measured at OD_550_ after 24h of growth in KA minimal medium. (B) Swim Zone (in millimeters) of the WT, Δ*fleQ,* Δ*rsmA*Δ*rsmE*Δ*rsmI*, and Δ*fleQ* Δ*rsmA*Δ*rsmE*Δ*rsmI* strains after toothpick inoculation on KA medium supplemented with 0.3% agar and 24h of growth at 30°C. (C) Quantification of cell surface-associated LapA after 16h of growth on KA agar as described in Figure 1. Representative images are included above each graph. Statistical significance for this figure was determined using one-way ANOVAs with Tukey’s multiple comparisons tests. *, P<0.05; **, P<0.01; ****, P<0.0001. All error bars represent standard deviation.

Given that the Gac/Rsm Pathway has been shown to regulate motility in various pseudomonads, we assessed these mutants for motility on KA medium supplemented with 0.3% agar. After 24h of growth, the Δ*rsmA*, Δ*rsmE*, and Δ*rsmI* single mutants show no significant difference in motility compared to the WT or Δ*fleQ* parental strains (Fig. S5B). Interestingly, the Δ*rsmA* Δ*rsmE* Δ*rsmI* triple mutant in the WT background showed a significant decrease in motility compared to the parental strain whereas the triple mutant in the Δ*fleQ* background was non-motile like the parental strain (Fig. 5B). The reason for the reduction in motility in the Δ*rsmA* Δ*rsmE* Δ*rsmI* triple mutant is not currently understood.

Given these biofilm and motility results, we then assessed LapA localization on the cell surface of the Δ*rsmE* single mutant and Δ*rsmA* Δ*rsmE* Δ*rsmI* triple mutant via dot blot analysis. After 16h of growth on KA agar, the Δ*rsmA* Δ*rsmE* Δ*rsmI* and Δ*rsmE* mutants showed a significant increase in LapA surface levels in the Δ*fleQ* background, but not the WT background (Fig. 5C and S5C).

Since the biofilm results suggest specificity amongst the Rsm proteins, we assessed the specificity of the small regulatory RNAs by overexpressing the RsmX, RsmY, or RsmZ sRNAs in a Δ*fleQ* mutant from the IPTG-inducible plasmid pMQ123 and assessed the strains for biofilm formation. After 24h of growth in KA medium supplemented with 1mM IPTG for induction, the Δ*fleQ* mutant expressing RsmZ showed a modest, but statistically significant increase in the biofilm formed compared to growth in KA alone (no inducer, Fig 6A).

**FIG 6.**
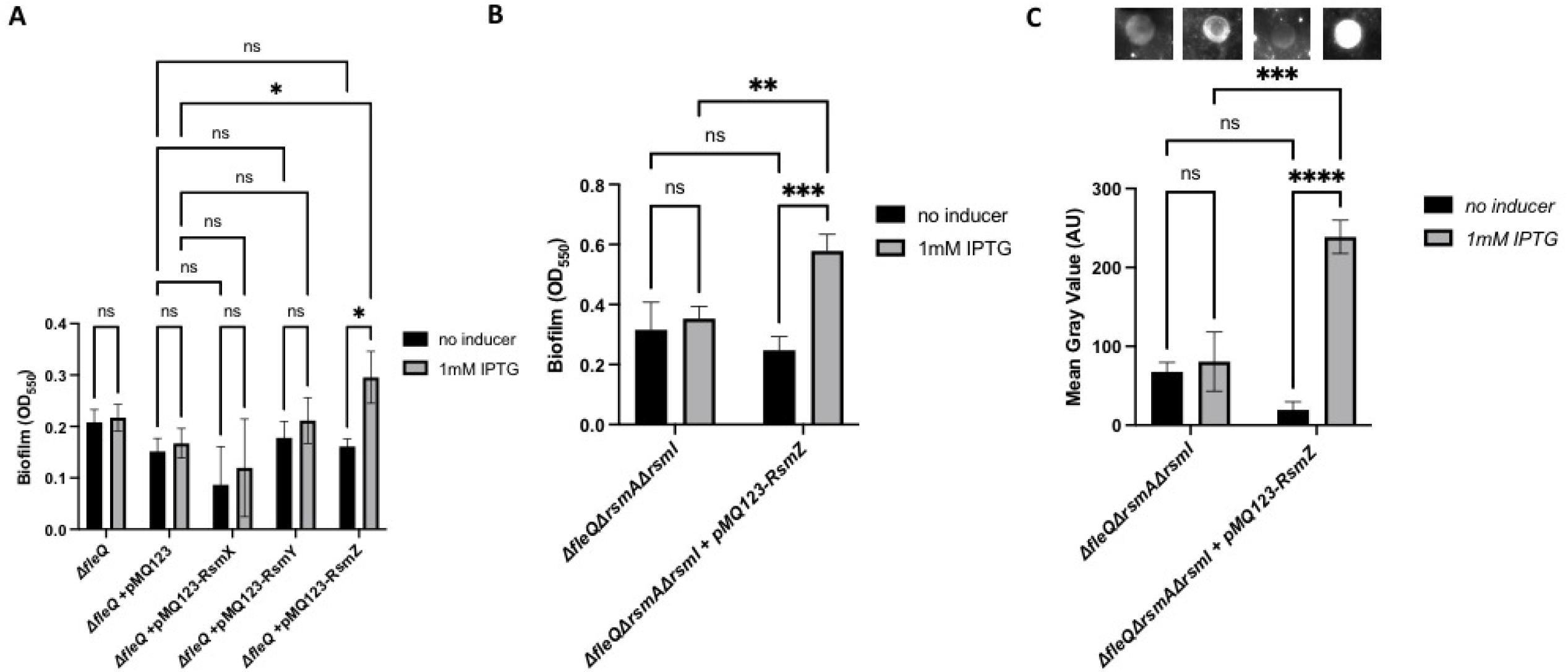
Overexpressing the small regulatory RNA RsmZ in a FleQ-deficient increases biofilm formation. (A) Quantification of the biofilm formed by the Δ*fleQ* strain with and without pMQ123, pMQ123-RsmX, pMQ123-RsmY, or pMQ123-RsmZ measured at OD_550_ after 24h of growth in KA minimal medium with or without 1mM IPTG for induction. (B) Quantification of the biofilm formed by the Δ*fleQ*Δ*rsmA*Δ*rsmI* strain with and without the pMQ123-RsmZ construct measured at OD_550_ after 24h of growth in KA minimal medium with or without 1mM IPTG for induction. (C) Quantification of cell surface-associated LapA after 16h of growth on KA agar with or without 1mM IPTG for induction as described in Figure 1. Statistical significance for this figure was determined using two-way ANOVAs with Tukey’s multiple comparisons tests. *, P<0.05; **, P<0.01; ***, P<0.001; ****, P<0.0001. All error bars represent standard deviation.

Given this finding, we then overexpressed RsmZ in a Δ*fleQ* Δ*rsmA* Δ*rsmI* mutant, which still has the ability to produce RsmE, and assessed the strain for biofilm formation. After 24h of growth in KA medium supplemented with 1mM IPTG for induction, the Δ*fleQ* Δ*rsmA* Δ*rsmI* mutant overexpressing RsmZ showed a significant increase in the biofilm formed compared to growth in KA alone (no inducer, Fig 6B). We then assessed these strains for cell surface levels of LapA via dot blot analysis. After 16h of growth on KA agar supplemented with 1mM IPTG, the Δ*fleQ* Δ*rsmA* Δ*rsmI* mutant overexpressing RsmZ showed a significant increase in LapA surface levels compared to growth on KA agar alone (no inducer, Fig. 6C).

### A mutation in the *gacA* gene does not impact early biofilm formation in KA medium

Previous work in *P. aeruginosa* demonstrated that the Gac/Rsm pathway regulates biofilm formation in both the PA01 and PA14 strains and that deletion of either *gacS* or *gacA*, which disrupts the Gac/Rsm pathway, leads to a large reduction in biofilm formation (38, 43). To assess the impact of disrupting the Gac/Rsm pathway in *P. fluorescens* Pf0-1, we made a chromosomal deletion of *gacA* in the WT and Δ*fleQ* strains and assessed these mutants for biofilm formation. Interestingly, after 24h of growth in KA medium, the Δ*gacA* mutant showed no difference in biofilm formation compared to the WT strain and the Δ*gacA*Δ*fleQ* mutant showed no difference in biofilm formation compared to the Δ*fleQ* mutant (Fig. 7A).

**FIG 7.**
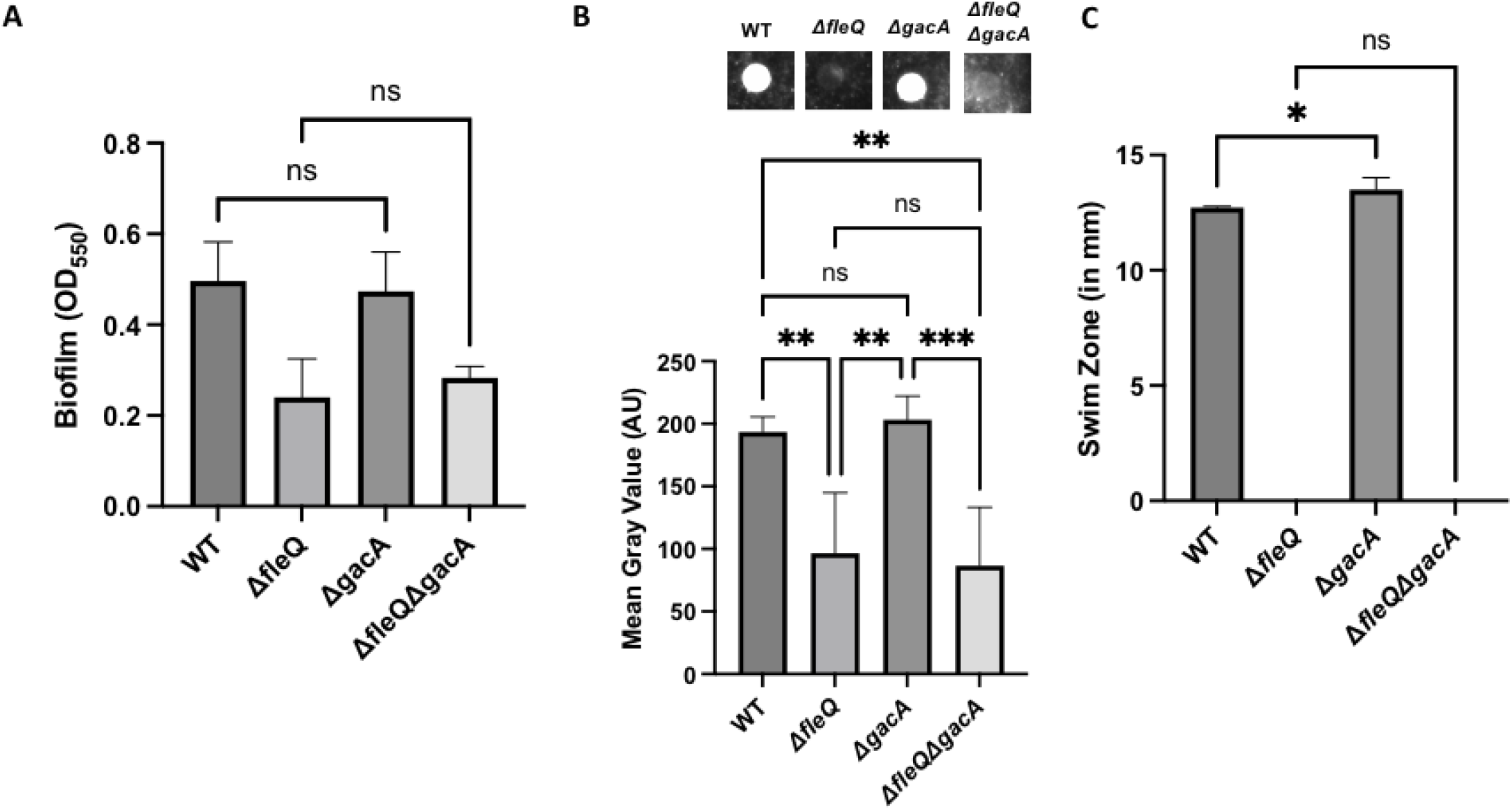
The Gac/Rsm system contributes to LapA production. (A) Quantification of the biofilm formed by the WT, Δ*fleQ,* Δ*gacA*, and Δ*fleQ*Δ*gacA* strains measured at OD_550_ after 24h of growth in KA minimal medium. (B) Quantification of cell surface-associated LapA after 16h of growth on KA agar as described in Figure 1. Statistical significance for this figure was determined using one-way ANOVAs with Tukey’s multiple comparisons tests. *, P<0.05; **, P<0.01; ***, P<0.001. All error bars represent standard deviation. (C) Swim Zone (in millimeters) of the WT, Δ*fleQ,* Δ*gacA*, and Δ*fleQ*Δ*gacA* strains after toothpick inoculation on KA supplemented with 0.3% agar and 24h of growth at 30°C.

The Δ*gacA* and Δ*gacA*Δ*fleQ* mutants were assessed for cell surface levels of LapA via dot blot analysis. After 16h of growth on KA agar, the Δ*gacA* mutant showed no significant difference in surface LapA compared to the WT strain, and the Δ*fleQ* Δ*gacA* mutant shows no significant difference in surface LapA compared to Δ*fleQ* strain (Fig. 7B).

Given that the Gac/Rsm pathway has been implicated in regulating motility, we next assessed the motility of these mutants on KA medium supplemented with 0.3% agar. The Δ*gacA* mutant showed a modest but significant increase in swimming motility compared to the WT strain, which is consistent with previous work in other pseudomonads. The Δ*fleQ* Δ*gacA* mutant also showed no significant difference in swimming motility from the parental Δ*fleQ* strain; both strains were non-motile (Fig. 7C). Together, these results suggest that the Gac/Rsm pathway is not strictly required for biofilm formation, but that activation of the pathway can enhance biofilm formation.

### Replacing the native *lapA* promoter with the constitutive P*_lac_* promoter partially overcomes FleQ-dependent regulation of LapA

Given that FleQ post-transcriptionally regulates LapA production, we posited that further increasing *lapA* expression by replacing the native *lapA* promoter with the non-native, constitutively active P*_lac_* promoter would partially rescue biofilm formation in a FleQ-deficient strain.

We inserted the constitutive P*_lac_* promoter in front of the *lapA* open reading frame in the WT and Δ*fleQ* strains, replacing the regulatory and the 5’ untranslated regions of the *lapA* gene, and assessed these strains for biofilm formation. After 24h of growth in KA medium, the strain carrying the P*_lac_* promoter in the Δ*fleQ* background showed significantly more biofilm formed compared to the parental strain (Fig. 8A). In contrast, the strain carrying the P*_lac_* promoter in WT background showed no difference in the biofilm formed compared to the parent strain (Fig. 7A).

**FIG 8.**
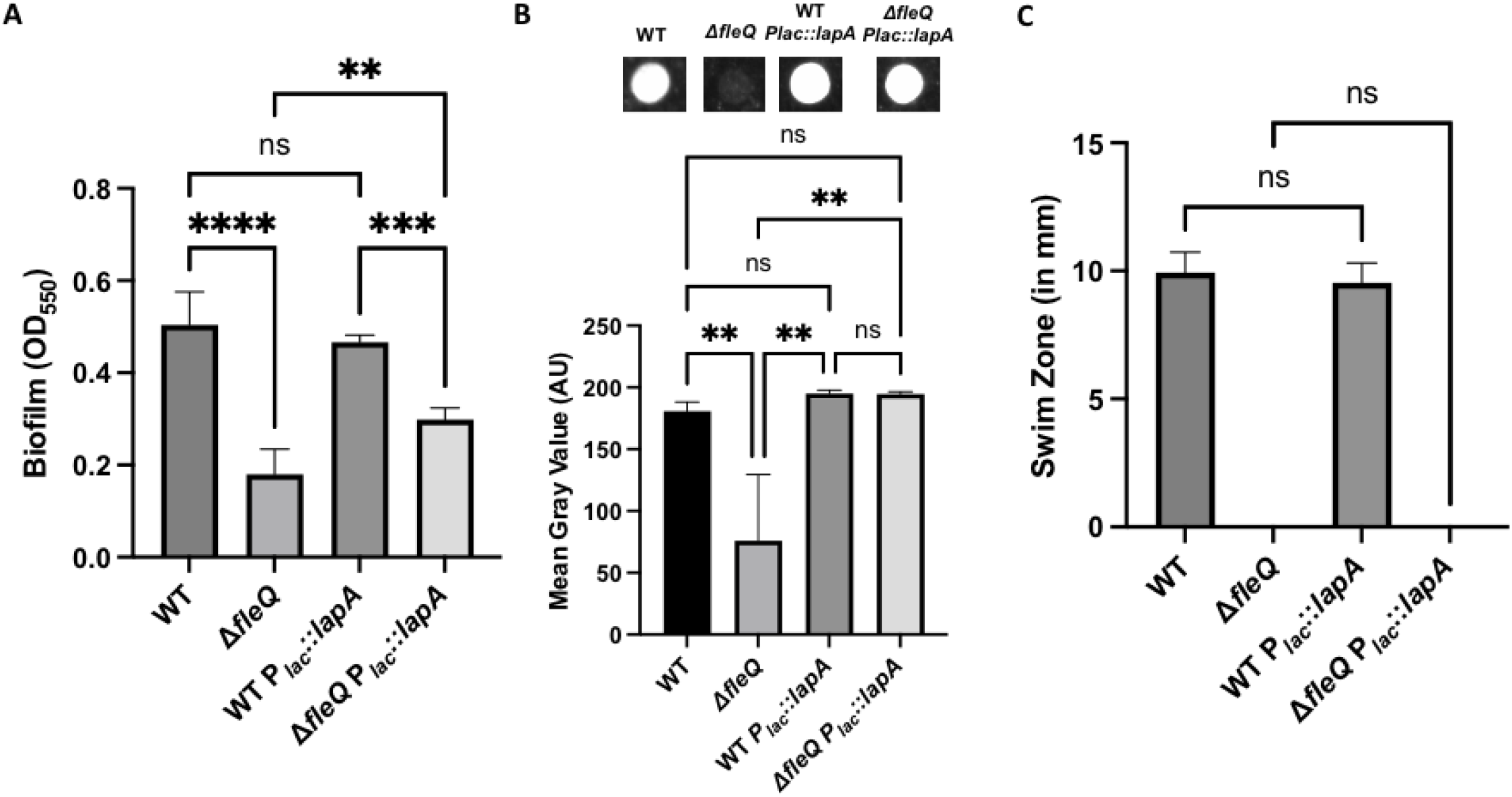
Replacing the native *lapA* promoter with a non-native FleQ-independent promoter partially restores biofilm formation in a FleQ-deficient strain. (A) Quantification of the biofilm formed by the WT and Δ*fleQ* strains with the native *lapA* or P*_lac_* promoter measured at OD_550_ after 24h of growth in KA minimal medium. (B) Quantification of cell surface-associated LapA after 16h of growth on KA agar as described in Figure 1. (C) Swim Zone (in millimeters) of the WT and Δ*fleQ* strains with the native *lapA* or P*_lac_* promoter strains after toothpick inoculation on KA supplemented with 0.3% agar and 24h of growth at 30°C. Statistical significance for this figure was determined using one-way ANOVAs with Tukey’s multiple comparisons tests. **, P<0.01; ***, P<0.001; ****, P<0.0001. All error bars represent standard deviation.

We next assessed the P*_lac_* promoter variants for LapA levels on the cell surface via dot blot analysis. After 16h of growth on KA agar, the WT P*_lac_* promoter variant showed no significant difference in cell surface-associated LapA as compared to the WT parental strain (Fig. 8B). In contrast, the Δ*fleQ* P*_lac_*::*lapA* strain showed significantly more cell surface-associated LapA as compared to the Δ*fleQ* parental strain. This increased LapA likely accounts for the increased biofilm formation observed for the Δ*fleQ* P*_lac_*::*lapA* strain. Collectively, these data indicate that sequences within the *lapA* promoter region are required to mediate FleQ-dependent regulation of this adhesin, a finding consistent with the phenotypes of the Box 2 mutants described above.

To confirm that driving *lapA* expression from the constitutive P*_lac_* promoter does not impact motility, we assessed these strains on motility agar. As expected, the WT carrying the P*_lac_* promoter swam like its parental strain, whereas the P*_lac_* promoter variant of the Δ*fleQ* background was non-motile, like the Δ*fleQ* mutant (Fig. 8C).

## Discussion

Our data show that FleQ post-transcriptionally regulates biofilm formation by *P. fluorescens*. Our genetic analyses show that the loss of FleQ leads to a decrease in biofilm formation, which is attributed, at least in part, to the reduction of LapA and MapA production, and thus the lack of these adhesins on the cell surface. Whereas FleQ-dependent regulation is typically thought of as being transcriptional (26, 28, 29), our qRT-PCR analysis reveal that transcription of *lapA* and *mapA* genes are increased in a FleQ-deficient strain compared to a WT strain despite a reduction in protein level, which suggests that FleQ-dependent regulation of these adhesins is predominantly post-transcriptional. However, our subsequent mutagenic study of the predicted FleQ binding boxes in the *lapA* promoter (29) along with the qRT-PCR data suggest that FleQ does have a transcriptional effect on the *lapA* gene, but this alone is not sufficient to explain the decrease in biofilm formation in the FleQ-deficient strain. Despite the loss of biofilm formation, additional genetic and biochemically analyses utilizing our knowledge of LapD-dependent biofilm formation (34) show that a basal amount of LapA and MapA is still produced in a FleQ-deficient strain, which is sufficient to sustain biofilm formation in LapG-deficient background. Interestingly, we did not observe restoration of biofilm formation to wild-type levels in the *lapG* mutant, which we attribute to the loss of motility in a FleQ-deficient strain; motility plays a key positive role in early biofilm formation by this organism (47).

Our transposon mutagenesis identified 4 candidates that were able to restore biofilm formation in a FleQ-deficient strain, which included the histidine-kinase GacS that has been previous implicated in post-transcriptional regulation. This mutant showed high levels of *gacS* gene expression, which suggests overstimulation of the Gac/Rsm pathway. We subsequently found that mimicking an overstimulation of this pathway by deleting all of the target downstream regulators RsmA, RsmE, and RsmI restored biofilm formation in a FleQ-deficient strain, much like the *gacS*::*Tn*M mutant from the transposon screen. Interestingly, when deleting these regulators individually, we only observed a restoration of biofilm formation when the *rsmE* gene is deleted. This observation is likely due to the nature of the Rsm proteins having distinct regulomes with a subset of overlapping targets, which has been previously demonstrated in *Pseudomonas putida* (48). We similarly stimulated the Gac/Rsm pathway by overexpressing the small regulatory RNAs and found that biofilm formation was only restored in a FleQ-deficient strain when RsmZ was expressed from a multi-copy plasmid. This specificity is likely due to the variability of binding affinities between the small regulatory RNAs and Rsm proteins, which has previously been demonstrated for *P. aeruginosa* (45). A recently published RNA-seq dataset revealed that the *gacA* gene is significantly downregulated in a FleQ-deficient strain compared to a wild-type strain (31), suggesting that the Gac/Rsm pathway activity is attenuated in a FleQ-deficient strain. This published finding along with our results implicate the Gac/Rsm pathway in the FleQ-dependent post-transcriptional regulation of biofilm formation. Interestingly, we show that a GacA-deficient strain, which would abolish signaling through the Gac/Rsm pathway, is not sufficient to reduce biofilm formation in a wildtype strain, suggesting additional regulatory mechanisms involved in biofilm formation that have yet to be elucidated.

## Materials and Methods

### Strains and media used in this study

The strains and plasmids used in this study are listed in **Table S1**. *P. fluorescens* Pf0-1 and *E. coli* S17-1 λ-pir, SM10 λ-pir, and JM109 were used throughout this study. *E. coli* was routinely grown in lysogeny broth (LB) and *P. fluorescens* was routinely grown on LB or on KA minimal medium, as previously defined by Collins et al. (18), containing 50 mM Tris-HCl (pH 7.4), 0.61 mM MgSO_4_, 1 mM K_2_HPO_4_, and 0.4% (wt/vol) L-arginine HCl, or KA supplemented with 1.5% agar. Medium was supplemented with 50 µg/ml carbenicillin for *E. coli* strains harboring the transposon-containing shuttle vector pBT20, miniTn*7*/pMQ56, or the Tn7 transposase-containing helper plasmid pTns3, or with 10 ug/ml gentamycin when harboring the allelic exchange plasmid pMQ30 or the expression plasmids pMQ72 and pMQ123. For *P. fluorescens*, the medium was supplemented with 150 µg/ml kanamycin when harboring pSMC21 or 30 µg/ml gentamycin when harboring pMQ72, pMQ123, or miniTn*7* integrated into the *att* site.

### Construction of in-frame chromosomal deletion mutants

For all chromosomal deletions, the allelic exchange vector pMQ30 was first digested with SmaI (New England BioLabs) and ∼1.5kb flanking regions of the targeted gene were amplified using Phusion polymerase (New England BioLabs) with primers containing 20bp of homology to the pMQ30 SmaI cut-site at their 5’ end. All primers used in the study are listed in **Table S2**. These amplicons were inserted into pMQ30 using the NEBuilder HiFi DNA Assembly kit (New England BioLabs) according to the manufacturer specifications, and the constructs were electroporated in *E. coli* S17-1 λ-pir. After recovery in LB for 1h at 30°C, cells were plated on LB agar with 10 µg/ml gentamycin; candidates were sequenced to confirm fragment insertion into the plasmid. Constructs were introduced into *P. fluorescens* through conjugation, whereby 1 mL of *P. fluorescens* and 1 mL of *E. coli* were mixed in a 2 mL microcentrifuge tube, pelleted, washed in LB, and then plated on LB agar supplemented with 30 µg/ml gentamycin and 30 µg/ml chloramphenicol to select for *P. fluorescens* merodiploids. Cells were then plated on LB agar without sodium chloride supplemented with 10% sucrose to facilitate looping out of the drug resistance cassette. Deletions were confirmed with PCR amplification and Sanger sequencing.

### Construction of *fleQ* complementation mutant

For *fleQ* complementation, the vector miniTn*7*/pMQ56 was first digested with SmaI (New England BioLabs) and the *fleQ* gene and its native promoter were amplified using Phusion polymerase (New England BioLabs) with primers containing 20bp of homology to the mTn7/pMQ56 SmaI cut-site at their 5’ end. All primers used in the study are listed in **Table S2**. The amplicon was inserted into miniTn*7*/pMQ56 using the NEBuilder HiFi DNA Assembly kit (New England BioLabs) according to the manufacturer specifications, and the constructs were electroporated in *E. coli* S17-1 λ-pir. After recovery in LB for 1h at 30°C, cells were plated on LB agar supplemented with 10µg/ml gentamycin; candidates were sequenced to confirm fragment insertion into the plasmid. Plasmid DNA was isolated using the QIAprep Spin Miniprep Kit (Qiagen) and introduced into *P. fluorescens* through conjugation with the construct and the helper plasmid pTns3 containing the miniTn*7* transposase machinery, whereby 500µl of the three strains was mixed in a 2 mL microcentrifuge tube. Cells were pelleted, washed in LB, and then plated on LB agar supplemented with 30 µg/ml gentamycin and 30 µg/ml chloramphenicol to select for miniTn*7* insertion into *P. fluorescens*. Incorporation onto the genome was confirmed via PCR amplification and Sanger sequencing.

### Construction of small regulatory RNA overexpression constructs

For all overexpression constructs, the allelic *lacI*-containing shuttle vector pMQ123 was first digested with BamHI (New England BioLabs) and the targeted small regulatory RNAs were amplified using Phusion polymerase (New England BioLabs) with primers containing 20bp of homology to the pMQ123 BamHI cut-site at their 5’ end. All primers used in the study are listed in **Table S2**. The amplicons were inserted into pMQ123 downstream of the P*tac* promoter using the NEBuilder HiFi DNA Assembly kit (New England BioLabs) according to the manufacturer specifications, and the constructs were electroporated in *E. coli* S17-1 λ-pir. After recovery in LB for 1h at 30°C, cells were plated on LB agar supplemented with 10 µg/ml gentamycin; candidates were sequenced to confirm fragment insertion into the plasmid. Plasmid DNA was isolated using the QIAprep Spin Miniprep Kit (Qiagen) and introduced into *P. fluorescens* through electroporation. Cells were allowed to recover in LB for 1h at 30°C and then were plated on LB agar supplemented with 30 µg/ml gentamycin to select for retention of the construct.

### Biofilm formation in static 96-well plate

*P. fluorescens* Pf0-1 strains were grown in 5 ml of LB at 30°C for ∼16 hours with agitation. 1.5 ul of inoculum was added to 100 µl of KA minimal medium in each well of a 96-well round-bottom polypropylene plate, and the plate was incubated for the indicated time at 30°C in a humidified chamber. After incubation, supernatants were discarded and wells were washed once in water. 125 µl of 0.1% (wt/vol) crystal violet was pipetted into each well and the plate was incubated for 45 min at room temperature. Wells were then washed twice in water to remove excess crystal violet, and then the plate was incubated at 37°C for 1h to dry the wells. 150µl of a 45% methanol, 45% water, and 10% glacial acetic acid solution (destain solution) was added to each well and the plate was incubated for 5 min at 25°C. 100µl was then transferred from each well to a 96-well flat-bottom polystyrene plate and measured on a spectrophotometer at an optical density of 550nm (OD_550_).

### Biofilm formation in microfluidic devices

Bacterial strains were grown in 5 ml LB supplemented with 150 ug/ul kanamycin to select for the pSMC21 plasmid overnight at 30 °C with agitation. 1 mL of overnight culture was transferred to a 1.5 mL microcentrifuge tube and the cells pelleted. Cell pellets were washed twice in KA medium and pellets resuspended in 100 µl of KA medium, and cultures were normalized to an OD_600_ of 1.5 in a 96 well plate.

Microfluidic devices were assembled and operated as previously described in Collins et al. (18), except in these studies the medium used was KA. After 120 hours of growth at room temperature, biofilms were imaged using a Nikon Eclipse Ti inverted microscope equipped with a Plan Fluor 40 DIC M N2 objective and Hamamatsu ORCA-Flash 4.0 camera. GFP was excited using a 488-nm laser and images were quantified using Fiji: ImageJ (49), as reported (50).

### Quantitative reverse transcription PCR

Bacterial strains were grown in 5 ml LB overnight at 30°C with agitation and then 400µl of inoculum was pipetted onto KA medium supplemented with 1.5% agar. Following incubation, cells were scraped into 1 mL fresh KA medium using disposable cell scrapers with 20 mm blade width (VWR). Cells were pelleted with the respective supernatants discarded and then flash frozen in a dry ice-ethanol bath to preserve RNA integrity and stored at −80°C. In preparation for RNA extraction, pellets were thawed and resuspended in 100µl of 3mg/ml lysozyme and incubated at room temperature for 3 minutes. RNA was then extracted and isolated from cells using the Zymo Direct-zol RNA miniprep kit with TRI Reagent according to the manufacturer specifications. The isolated RNA was then treated twice with the TURBO DNA-*free* kit (Invitrogen) according to manufacturer specifications, to remove contaminating genomic DNA from the samples. The isolated RNA was used to generate complementary DNA using the RevertAid First Strand cDNA Synthesis Kit (Thermo Scientifc) according to manufacturer specifications. Quantitative PCR was conducted using the complementary DNA and SsoFast EvaGreen Supermix (Bio-Rad) using the manufacturer specifications for reaction preparation, primer design, and thermocycler profile, and reactions were run in a CFX 96 Touch Real-Time PCR Detection System (Bio-Rad). All primers used for quantitative PCR are listed in **Table S2**.

### Transposon mutagenesis

*P. fluorescens* Pf0-1 Δ*fleQ* mutant strain and *E. coli* JM109 strain carrying the pBT20 plasmid were grown in LB (no antibiotics) for *P. fluorescens* and 50 µg/ml carbenicillin for *Ec* overnight at 30°C, with agitation. 1mL of each culture was mixed in a 1.5 mL microcentrifuge tube and centrifuged for 3 minutes at 13,000 rpm. Cell pellets were washed in fresh LB and then resuspended in 100 µl of LB. The mixture was plated on LB agar to facilitate conjugation of the pBT20 plasmid into *P. fluorescens*. Plates were incubated for 90 min at 30°C, and the cell mixture was scraped up in 1 mL LB with a disposable cell scraper with 20 mm blade width. The cell pellet was washed once in fresh LB, and then dilutions were plated on LB agar supplemented with 30 µg/ml gentamycin and 30 ug/ml chloramphenicol to select for *P. fluorescens* mutants with chromosomal transposon insertions and against the *E. coli* donor, respectively. Individual candidates were picked with sterile pipette tips and inoculated in LB in sterile 96-well flat-bottom polystyrene plates and the plates were incubated at 30°C for 16 h. 100 µl of 10% sterile glycerol was added to each well and plates were frozen at −80°C. For every independent replicate of a conjugation, several random wells were chosen for arbitrary primed PCR analysis (see below) to verify the presence of *P. fluorescens* Pf0-1 containing a transposon. Mutants were then screened for their ability to form biofilm by using a 96-pin replicator (Dankar) to transfer inoculum from the frozen plates to 96-well round-bottom polypropylene plates (Corning) containing 100 µl of fresh KA medium. Plates were incubated and wells quantified according to the biofilm formation CV assay described above. Candidates with increased biofilm formation compared to the Δ*fleQ* mutant parental strain were struck out for single colonies from the freezer plate stock onto fresh LB agar with 30 µg/ml gentamycin and incubated overnight at 30°C. A single colony was picked with a sterile pipette tip and used to inoculate LB supplemented with 30 µg/ml gentamycin, which was incubated overnight at 30°C with agitation. 750 µl of inoculum and 750 µl of sterile 10% glycerol were mixed in 2 mL cryovials (Nunc) and stored at −80°C. From these frozen stocks, candidates were rescreened for biofilm formation in KA medium for 24h according to the biofilm formation method described above. For candidates with increased biofilm formation compared to the Δ*fleQ* mutant parental strain, transposon chromosomal insertion sites were determined using arbitrary primed PCR as described by O’Toole et al. (47). Based on these results, primers were designed to amplify and sequence the chromosomal flanks of the predicted insertion site to confirm the exact insertion location.

### Swim assay

*P. fluorescens* Pf0-1 strains were initially grown overnight in 5 ml of LB at 30°C hours with agitation, and then 1mL aliquots were transferred to 1.5 mL microcentrifuge tubes. Sterile toothpicks were used to stab inoculate KA supplemented with 0.3% agar (KA Swim Agar), and plates were incubated for 24 hours at 30°C. The diameter of resulting swim zones was measured using a ruler.

### Lap and MapA cell surface levels via dot blot analysis

*P. fluorescens* Pf0-1 strains were initially grown in 5 ml of LB at 30°C for ∼16 hours with agitation, and then 400 ul of inoculum was pipetted onto KA medium supplemented with 1.5% agar. The plates were incubated at 30°C for either 16h or 24h, for LapA-HA- and MapA-HA-tagged strains, respectively. Following incubation, cells were scraped into 1mL fresh KA medium using disposable cell scrapers with 20 mm blade width (VWR). Cell pellets were washed once and then resuspended in 110 µl of fresh KA medium. In a 96 well plate, cultures were normalized and diluted to a final OD_600_ value of 10. 10 µl of normalized cell culture was spotted on 0.2 µm pore-size nitrocellulose membrane (Bio-Rad) and allowed to dry. The membrane was then incubated in Tris-Buffered Saline (TBS; Bio-Rad) supplemented with 3% BSA (Sigma) and 0.1% Tween 20 (TBST; Sigma) for 1 hour at 25°C and then transferred to TBST supplemented with purified anti-HA antibody clone 16B12 (Biolegend) at a 1:2000 dilution for 1h at 25°C. The nitrocellulose membrane was washed three times in TBST for 5 minutes at 25°C, and then transferred to TBST supplemented with rabbit anti-mouse IgG antibody conjugated to horseradish peroxidase (Bio-Rad) at a 1:15000 dilution for 1h at 25°C. The membrane was then washed 3 times in TBST for 5 minutes at 25°C and then once in TBS for 5 minutes at 25°C to wash away excess Tween 20. The nitrocellulose was overlaid with a mixture of 1mL enhanced luminol and 1mL oxidizing reagent for 30s at 25°C per specification of the Western Lightning ECL Pro Kit (Perkin-Elmer), and then the blots were exposed on a BioRad ChemiDoc MP Imaging System. Images were imported to ImageJ (NIH) and the dots were quantified as previously described in Collins et al. (18).

### Whole cell lysate Western blot analysis

*P. fluorescens* Pf0-1 strains were initially grown in 5 ml of LB at 30°C for ∼16 hours with agitation. For LapA-HA tagged strains, 400 ul of inoculum was pipetted onto KA medium supplemented with 1.5% agar and incubated at 30°C for 16h. Following incubation, cells were scraped into 1mL fresh KA medium using disposable cell scrapers with 20mm blade width (VWR). For MapA-HA tagged strains, 1mL of inoculum was pipetted into 25 mL of KA medium and incubated at 30°C for 24h with agitation at 200rpm. After incubation, cultures were transferred to 50 mL conical centrifuge tubes. Cells were centrifuged and then resuspended in 100 µl of 3 mg/mL lysozyme and incubated at room temperature for 5 minutes. Reactions were incubated on ice and sonicated four times for 5 seconds at 30% amplitude, with incubation on ice between replicates. Cell slurries were spun down at 4°C and cell lysates were transferred to fresh 1.5 mL centrifuge tubes. Serial dilutions of cell lysates were made, and protein was quantified using the Pierce BCA Protein Assay Kit (ThermoFisher) according to the manufacturer specifications. 25 µg or 50 µg of normalized protein for LapA-HA or MapA-HA tagged strains respectively was mixed with 4x Laemmli Sample Buffer (Bio-Rad) and 2-mercaptoethanol to a 1x working dilution and samples were boiled for 5 minutes. Samples were loaded in 7.5% Mini-Protean TGX precast gels (Bio-Rad) with either 3 ul HiMark Pre-Stained Protein Standard (Thermo Fisher) for LapA or 3 ul Spectra Multicolor High Range Protein Ladder (Thermo Fisher) for MapA, and resolved for 4 hours at 120V for LapA and 3 hours at 115V for MapA in Mini-PROTEAN Tetra Vertical Electrophoresis Cells (Bio-Rad) with 1x Tris/Glycine/SDS buffer (Bio-Rad). The gel was transferred to 0.2 µm pore-size nitrocellulose membrane (Bio-Rad) using the Transblot Turbo Transfer System (Bio-Rad) according to manufacturer specifications, using the high molecular weight setting. The membrane was then incubated in Tris-Buffered Saline (TBS; Bio-Rad) supplemented with 3% BSA (Sigma) and 0.1% Tween 20 (TBST; Sigma) for 1h at 25°C and then transferred to TBST supplemented with purified anti-HA antibody clone 16B12 (Biolegend) at a 1:2000 dilution for 1h at 25°C. The nitrocellulose membrane was washed three times in TBST for 5 minutes at 25°C, and then transferred to TBST supplemented with rabbit anti-mouse IgG antibody conjugated to horseradish peroxidase (Bio-Rad) at a 1:15000 dilution for 1h at 25°C. The membrane was then washed 3 times in TBST for 5 minutes at 25°C and then once in TBS for 5 minutes at 25°C to wash away excess Tween 20. The nitrocellulose was overlaid with a mixture of 1mL enhanced luminol and 1mL oxidizing reagent for 30s at 25°C per specification of the Western Lightning ECL Pro Kit (Perkin-Elmer), and then the blots were exposed on a BioRad ChemiDoc MP Imaging System. Images were imported to ImageJ (NIH) and blots were quantified by defining a region of interest (ROI) and using that object to measure each band for mean gray value. The background signal was determined by measuring an area of the blot without a band and subtracting this value from the measured bands.

### Data availability

This manuscript has no large datasets.

## Acknowledgement

We would like to thank Dr. Shanice Webster and Dr. Sherry Kuchma for numerous helpful discussions. We would especially like to thank Dr. Fabrice Jean-Pierre for many insightful conversations regarding the Gac/Rsm system in multiple *Pseudomonas* spp. We thank Dr. Carey Nadell and his laboratory for their generous gift of microfluidics devices. Sequencing services were provided by the Dartmouth Molecular Biology Core. This work was supported by NIH grants R01GM123609 and R01AI168017 to G.A.O. The Molecular Biology Core was supported by NIH grant P30CA023108.

## Literature Cited.

1. Paulsen IT, Press CM, Ravel J, Kobayashi DY, Myers GSA, Mavrodi DV, DeBoy RT, Seshadri R, Ren Q, Madupu R, Dodson RJ, Durkin AS, Brinkac LM, Daugherty SC, Sullivan SA, Rosovitz MJ, Gwinn ML, Zhou L, Schneider DJ, Cartinhour SW, Nelson WC, Weidman J, Watkins K, Tran K, Khouri H, Pierson EA, Pierson LS, Thomashow LS, Loper JE. 2005. Complete genome sequence of the plant commensal *Pseudomonas fluorescens* Pf-5. Nat Biotechnol 23:873–878.

2. Howell CR. 1979. Control of *Rhizoctonia solani* on cotton seedlings with *Pseudomonas fluorescens* and with an antibiotic produced by the bacterium. J Phytopathol 69:480.

3. Keel C. 1992. Suppression of root diseases by *Pseudomonas fluorescens* CHA0: importance of the bacterial secondary metabolite 2,4-diacetylphloroglucinol. MPMI 5:4.

4. Haas D, Défago G. 2005. Biological control of soil-borne pathogens by fluorescent pseudomonads. Nat Rev Microbiol 3:307–319.

5. Khabbaz RF, Arnow PM, Highsmith AK, Herwaldt LA, Chou T, Jarvis WR, Lerche NW, Allen JR. 1984. *Pseudomonas fluorescens* bacteremia from blood transfusion. Am J Med 76:62–68.

6. Hsueh P-R, Teng L-J, Pan H-J, Chen Y-C, Sun C-C, Ho S-W, Luh K-T. 1998. Outbreak of *Pseudomonas fluorescens* bacteremia among oncology patients. J Clin Microbiol 36:2914– 2917.

7. Gershman MD, Kennedy DJ, Noble-Wang J, Kim C, Gullion J, Kacica M, Jensen B, Pascoe N, Saiman L, McHale J, Wilkins M, Schoonmaker-Bopp D, Clayton J, Arduino M, Srinivasan A. 2008. Multistate outbreak of *Pseudomonas fluorescens* bloodstream infection after exposure to contaminated heparinized saline flush prepared by a compounding pharmacy. Clin Infect Dis 47:1372–1379.

8. Wong V, Levi K, Baddal B, Turton J, Boswell TC. 2011. Spread of *Pseudomonas fluorescens* due to contaminated drinking water in a bone marrow transplant unit. J Clin Microbiol 49:2093–2096.

9. Benito N, Mirelis B, Luz Gálvez M, Vila M, López-Contreras J, Cotura A, Pomar V, March F, Navarro F, Coll P, Gurguí M. 2012. Outbreak of *Pseudomonas fluorescens* bloodstream infection in a coronary care unit. J Hosp Infect 82:286–289.

10. Landers CJ, Cohavy O, Misra R, Yang H, Lin Y, Braun J, Targan SR. 2002. Selected loss of tolerance evidenced by Crohn’s disease–associated immune responses to auto- and microbial antigens. Gastroenterology 123:689–699.

11. Wei B, Huang T, Dalwadi H, Sutton CL, Bruckner D, Braun J. 2002. *Pseudomonas fluorescens* encodes the crohn’s disease-associated I2 sequence and T-cell superantigen. Infect Immun 70:6567–6575.

12. Madi A, Svinareff P, Orange N, Feuilloley MG, Connil N. 2010. *Pseudomonas fluorescens* alters epithelial permeability and translocates across Caco-2/TC7 intestinal cells. Gut Pathog 2:16.

13. Madi A, Lakhdari O, Blottière HM, Guyard-Nicodème M, Le Roux K, Groboillot A, Svinareff P, Doré J, Orange N, Feuilloley MG, Connil N. 2010. The clinical *Pseudomonas fluorescens* MFN1032 strain exerts a cytotoxic effect on epithelial intestinal cells and induces Interleukin-8 via the AP-1 signaling pathway. BMC Microbiol 10:215.

14. Alnabhani Z, Montcuquet N, Biaggini K, Dussaillant M, Roy M, Ogier-Denis E, Madi A, Jallane A, Feuilloley M, Hugot J-P, Connil N, Barreau F. 2015. *Pseudomonas fluorescens* alters the intestinal barrier function by modulating IL-1β expression through hematopoietic NOD2 signaling. Inflamm Bowel Dis 21:543–555.

15. Sauer K, Camper AK, Ehrlich GD, Costerton JW, Davies DG. 2002. *Pseudomonas aeruginosa* displays multiple phenotypes during development as a biofilm. J Bacteriol 184:1140–1154.

16. Sauer K, Stoodley P, Goeres DM, Hall-Stoodley L, Burmølle M, Stewart PS, Bjarnsholt T. 2022. The biofilm life cycle: expanding the conceptual model of biofilm formation. 10. Nat Rev Microbiol 20:608–620.

17. Hinsa SM, Espinosa-Urgel M, Ramos JL, O’Toole GA. 2003. Transition from reversible to irreversible attachment during biofilm formation by *Pseudomonas fluorescens* WCS365 requires an ABC transporter and a large secreted protein. Mol Microbiol 49:905–918.

18. Collins AJ, Pastora AB, Smith TJ, O’Toole GA. 2020. MapA, a second large RTX adhesin conserved across the pseudomonads, contributes to biofilm formation by *Pseudomonas fluorescens*. J Bacteriol 202:e00277–20.

19. Collins AJ, Smith TJ, Sondermann H, O’Toole GA. 2020. From input to output: The Lap/c-di-GMP biofilm regulatory circuit. Annu Rev Microbiol 74:607–631.

20. Monds RD, Newell PD, Schwartzman JA, O’Toole GA. 2006. Conservation of the Pho regulon in P*seudomonas fluorescens* Pf0-1. Appl Environ Microbiol 72:1910–1924.

21. Monds RD, Newell PD, Gross RH, O’Toole GA. 2007. Phosphate-dependent modulation of c-di-GMP levels regulates *Pseudomonas fluorescens* Pf0-1 biofilm formation by controlling secretion of the adhesin LapA. Mol Microbiol 63:656–679.

22. Giacalone D, Smith TJ, Collins AJ, Sondermann H, Koziol LJ, O’Toole GA. 2018. Ligand-mediated biofilm formation via enhanced physical interaction between a diguanylate cyclase and its receptor, mBio 9:13.

23. Dahlstrom KM, Collins AJ, Hogan DA, O’Toole GA. 2018. A multimodal strategy used by a large c-di-GMP network. J Bacteriol 200:19.

24. Pérez-Mendoza D, Felipe A, Ferreiro MD, Sanjuán J, Gallegos MT. 2019. AmrZ and FleQ Co-regulate cellulose production in *Pseudomonas syringae* pv. Tomato DC3000. Front Microbiol 10:746.

25. Molina-Henares MA, Ramos-González MI, Daddaoua A, Fernández-Escamilla AM, Espinosa-Urgel M. 2017. FleQ of *Pseudomonas putida* KT2440 is a multimeric cyclic diguanylate binding protein that differentially regulates expression of biofilm matrix components. Res Microbiol 168:36–45.

26. Banerjee P, Chanchal, Jain D. 2019. Sensor I regulated ATPase activity of FleQ is essential for motility to biofilm transition in *Pseudomonas aeruginosa*. ACS Chem Biol 14:1515– 1527.

27. Hickman JW, Harwood CS. 2008. Identification of FleQ from *Pseudomonas aeruginosa* as a c-di-GMP-responsive transcription factor. Mol Microbiol 69:376–389.

28. Baraquet C, Murakami K, Parsek MR, Harwood CS. 2012. The FleQ protein from *Pseudomonas aeruginosa* functions as both a repressor and an activator to control gene expression from the pel operon promoter in response to c-di-GMP. Nucleic Acids Res 40:7207–7218.

29. Baraquet C, Harwood CS. 2015. FleQ DNA binding consensus sequence revealed by studies of FleQ-dependent regulation of biofilm gene expression in *Pseudomonas aeruginosa*. J Bacteriol 198:178–186.

30. Blanco-Romero E, Redondo-Nieto M, Martínez-Granero F, Garrido-Sanz D, Ramos-González MI, Martín M, Rivilla R. 2018. Genome-wide analysis of the FleQ direct regulon in *Pseudomonas fluorescens* F113 and *Pseudomonas putida* KT2440. Sci Rep 8:13145.

31. Blanco-Romero E, Durán D, Garrido-Sanz D, Rivilla R, Martín M, Redondo-Nieto M 2022. Transcriptomic analysis of *Pseudomonas ogarae* F113 reveals the antagonistic roles of AmrZ and FleQ during rhizosphere adaption. Microb Genom 8:000750.

32. Garrido-Sanz D, Redondo-Nieto M, Martin M, Rivilla R 2021. Comparative genomics of the *Pseudomonas corrugata* subgroup reveals high species diversity and allows the description of *Pseudomonas ogarae* sp. nov. Microb Genom 7:000593.

33. Robleto EA, López-Hernández I, Silby MW, Levy SB. 2003. Genetic analysis of the AdnA regulon in *Pseudomonas fluorescens*: Nonessential role of flagella in adhesion to sand and biofilm formation. J Bacteriol 185:453–460.

34. Boyd CD, Chatterjee D, Sondermann H, O’Toole GA. 2012. LapG, required for modulating biofilm formation by *Pseudomonas fluorescens* Pf0-1, is a calcium-dependent protease. J Bacteriol 194:4406–4414.

35. Boyd CD, Smith TJ, El-Kirat-Chatel S, Newell PD, Dufrene YF, O’Toole GA. 2014. Structural features of the *Pseudomonas fluorescens* biofilm adhesin LapA required for LapG-dependent cleavage, biofilm formation, and cell surface localization. J Bacteriol 196:2775–2788.

36. Smith TJ, Font ME, Kelly CM, Sondermann H, O’Toole GA. 2018. An N-Terminal retention module anchors the giant adhesin LapA of *Pseudomonas fluorescens* at the cell surface: a novel subfamily of Type I secretion systems. J Bacteriol 200:e00734–17.

37. Monds RD, Newell PD, Wagner JC, Schwartzman JA, Lu W, Rabinowitz JD, O’Toole GA. 2010. Di-adenosine tetraphosphate (Ap4A) metabolism impacts biofilm formation by *Pseudomonas fluorescens* via modulation of c-di-GMP-dependent pathways. J Bacteriol 192:3011–3023.

38. Brencic A, McFarland KA, McManus HR, Castang S, Mogno I, Dove SL, Lory S. 2009. The GacS/GacA signal transduction system of *Pseudomonas aeruginosa* acts exclusively through its control over the transcription of the RsmY and RsmZ regulatory small RNAs. Mol Microbiol 73:434–445.

39. Heeb S, Blumer C, Haas D. 2002. Regulatory RNA as mediator in GacA/RsmA-dependent global control of exoproduct formation in *Pseudomonas fluorescens* CHA0. J Bacteriol 184:1046–1056.

40. Humair B, Wackwitz B, Haas D. 2010. GacA-controlled activation of promoters for small RNA genes in *Pseudomonas fluorescens*. Appl Environ Microbiol 76:1497–1506.

41. Valverde C, Heeb S, Keel C, Haas D. 2003. RsmY, a small regulatory RNA, is required in concert with RsmZ for GacA-dependent expression of biocontrol traits in *Pseudomonas fluorescens* CHA0. Mol Microbiol 50:1361–1379.

42. Kay E, Dubuis C, Haas D. 2005. Three small RNAs jointly ensure secondary metabolism and biocontrol in *Pseudomonas fluorescens* CHA0. Proc Natl Acad Sci U S A 102:17136– 17141.

43. Davies JA, Harrison JJ, Marques LLR, Foglia GR, Stremick CA, Storey DG, Turner RJ, Olson ME, Ceri H. 2007. The GacS sensor kinase controls phenotypic reversion of small colony variants isolated from biofilms of *Pseudomonas aeruginosa* PA14. FEMS Microbiol Ecol 59:32–46.

44. Reimmann C, Valverde C, Kay E, Haas D. 2005. Posttranscriptional repression of GacS/GacA-controlled genes by the RNA-binding protein RsmE acting together with RsmA in the biocontrol strain *Pseudomonas fluorescens* CHA0. J Bacteriol 187:276–85.

45. Janssen KH, Diaz MR, Golden M, Graham JW, Sanders W, Wolfgang MC, Yahr TL. 2018. Functional analyses of the RsmY and RsmZ small noncoding regulatory RNAs in *Pseudomonas aeruginosa*. J Bacteriol 200:e00736–17.

46. Sorger-Domenigg T, Sonnleitner E, Kaberdin VR, Bläsi U. 2007. Distinct and overlapping binding sites of *Pseudomonas aeruginosa* Hfq and RsmA proteins on the non-coding RNA RsmY. Biochem Biophys Res Commun 352:769–773.

47. O’Toole GA, Kolter R. 1998. Initiation of biofilm formation in P*seudomonas fluorescens* WCS365 proceeds via multiple, convergent signalling pathways: a genetic analysis. Mol Microbiol 28:449–461.

48. Huertas-Rosales Ó, Romero M, Chan K-G, Hong K-W, Cámara M, Heeb S, Barrientos-Moreno L, Molina-Henares MA, Travieso ML, Ramos-González MI, Espinosa-Urgel M. 2021. Genome-wide analysis of targets for post-transcriptional regulation by Rsm proteins in *Pseudomonas putida*. Front Mol Biosci 8:624061.

49. Schindelin J, Arganda-Carreras I, Frise E, Kaynig V, Longair M, Pietzsch T, Preibisch S, Rueden C, Saalfeld S, Schmid B, Tinevez J-Y, White DJ, Hartenstein V, Eliceiri K, Tomancak P, Cardona A. 2012. Fiji: an open-source platform for biological-image analysis. 7. Nat Methods 9:676–682.

50. Katharios-Lanwermeyer S, Koval SA, Barrack KE, O’Toole GA. 2022. The diguanylate cyclase YfiN of *Pseudomonas aeruginosa* regulates biofilm maintenance in response to peroxide. J Bacteriol 204:e00396–21.

